# Hydrogens and hydrogen-bond networks in macromolecular MicroED data

**DOI:** 10.1101/2022.04.08.487606

**Authors:** Max T.B. Clabbers, Michael W. Martynowycz, Johan Hattne, Tamir Gonen

## Abstract

Microcrystal electron diffraction (MicroED) is a powerful technique utilizing electron cryo-microscopy (cryo-EM) for protein structure determination of crystalline samples too small for X-ray crystallography. Electrons interact with the electrostatic potential of the sample, which means that scattered electrons carry informing about the charged state of atoms and can provide strong contrast for visualizing hydrogen atoms. Accurately identifying the positions of hydrogen atoms, and by extension the hydrogen bonding networks, is of importance for drug discovery and electron microscopy can enable such visualization. Using subatomic resolution MicroED data obtained from triclinic hen egg-white lysozyme, we identified hundreds of individual hydrogen atom positions and directly visualize hydrogen bonding interactions and the charged states of residues. Over a third of all hydrogen atoms are identified from strong difference peaks, the most complete view of a macromolecular hydrogen network visualized by electron diffraction to date. These results show that MicroED can provide accurate structural information on hydrogen atoms and non-covalent hydrogen bonding interactions in macromolecules. Furthermore, we find that the hydrogen bond lengths are more accurately described by the inter-nuclei distances than the centers of mass of the corresponding electron clouds. We anticipate that MicroED, coupled with ongoing advances in data collection and refinement, can open further avenues for structural biology by uncovering and understanding the hydrogen bonding interactions underlying protein structure and function.

## Main

Microcrystal electron diffraction (MicroED) has been successful in structure determination of crystalline biological specimens using electron cryo-microscopy (cryo-EM) (Nannenga, Shi, Leslie *et al*., 2014; Nannenga, Shi, Hattne *et al*., 2014; Yonekura *et al*., 2015), including novel structures (Rodriguez *et al*., 2015; Sawaya *et al*., 2016; Xu *et al*., 2019; Clabbers *et al*., 2021), as well as difficult to crystallize membrane proteins in detergents and lipids (Liu & Gonen, 2018; Martynowycz *et al*., 2020; Martynowycz, Shiriaeva *et al*., 2021). As electrons interact more strongly with matter than X-rays (Henderson, 1995), the crystal volume required for useful diffraction is typically about a million times smaller. Electrons are scattered by the electrostatic potential and the strength of scattering depends on the charged state of atoms (Cowley, 1995). The effects of charge distribution are already apparent at moderate to low resolution (Yonekura *et al*., 2015, 2018), and the charged state of residues in macromolecules has previously been investigated using electron crystallography (Kimura *et al*., 1997; Mitsuoka *et al*., 1999; Yonekura *et al*., 2015).

Electrostatic potential maps obtained from electron scattering can provide strong contrast for identifying hydrogen atoms, which has enabled localizing hydrogens in electron diffraction structures of small molecule organics and peptide fragments (Rodriguez *et al*., 2015; Sawaya *et al*., 2016; Dorset, 1995; Palatinus *et al*., 2017; Gruene *et al*., 2018; Jones *et al*., 2018; Clabbers *et al*., 2019; Takaba *et al*., 2021). Identifying the positions of hydrogen atoms and visualizing their resulting hydrogen bonding networks are crucial for understanding protein structure and function such as resolving precise drug or ligand binding interactions (Purdy *et al*., 2018; Clabbers *et al*., 2020; Martynowycz, Shiriaeva *et al*., 2021) or elucidating mechanisms for substrate transfer in membrane protein transporters and channels (Gonen *et al*., 2005; Liu & Gonen, 2018). In single-particle cryo-EM imaging, individual hydrogen atom positions were localized from reconstructions of apoferritin at 1.2 Å resolution (Nakane *et al*., 2020; Maki-Yonekura *et al*., 2021) and for the GABA_A_ receptor at 1.7 Å resolution (Nakane *et al*., 2020). Here, hydrogen atoms were identified by omitting them from the model and inspecting the peaks in a calculated *F_o_* – *F_C_* difference map following refinement in *Servalcat* based on crystallographic refinement routines implemented in *REFMAC5* (Murshudov *et al*., 2011; Yamashita *et al*., 2021). Since resolution is a local feature in cryo-EM, the accuracy of hydrogen identification varies across the map.

Visualizing hydrogen atoms in macromolecular crystallography generally requires (sub-) atomic resolution data, and the accuracy of localizing hydrogens varies with local structural flexibility which is reflected by the temperature factors. Typically, crystals of macromolecules are more disordered than peptides or small molecules and have a much higher solvent content. Therefore, in absence of atomic resolution data, identification of hydrogen atoms in macromolecular MicroED structures has remained elusive.

## Identifying hydrogen atoms in MicroED data

Recently, we reported the structure of triclinic hen egg-white lysozyme at 0.87 Å resolution using electron-counted MicroED data (Martynowycz, Clabbers, Hattne *et al*., 2021). MicroED data were collected from 16 crystal lamellae and the structure was phased *ab initio* as described previously (Supplementary Fig. 1) (Martynowycz, Clabbers, Hattne *et al*., 2021). Following density modification individual atoms could be resolved at sub-Ångström resolution, enabling automated model building of the entire structure without reference to a previously determined homologous model (Martynowycz, Clabbers, Hattne *et al*., 2021). The improvement in data accuracy and resolution in this study compared to previous efforts was realized by combining focused ion-beam milling to produce approximately 300 nm thin crystalline lamellae ideal for cryo-EM at 300 kV (Martynowycz, Clabbers, Unge *et al*., 2021), and collecting data in electron-counting mode at a significantly reduced exposure of only 0.64 e^-^.Å^-2^ per crystal dataset (Martynowycz, Clabbers, Hattne *et al*., 2021). A low exposure rate is required for electron counting as it ensures that the rate of scattered electrons remains within the linear range of the camera. Lowering the total exposure also reduces the effects of radiation damage that can affect the structural integrity of the protein and the ability to localize hydrogen atoms (Hattne *et al*., 2018; Leapman & Sun, 1995).

We set out to further refine the *ab initio* model resulting from automated building against the subatomic resolution MicroED data to closely examine individual hydrogen atom positions. First, the structural model was refined using electron scattering factors, isotropic atomic displacement parameters, and the default riding hydrogen model in *REFMAC5* (Murshudov *et al*., 2011). Twelve alternate side-chain conformations were modeled upon visual inspection using *Coot* (Emsley *et al*., 2010), and their occupancies were refined. The model was then refined using anisotropic *B*-factors until convergence (Supplementary Table 1). A crystallographic *mF_O_* – *DF_C_* difference map was calculated using a model without hydrogen atoms (Yamashita *et al*., 2021). Peaks in the difference hydrogen omit map at ≥2.0σ were then identified using *PEAKMAX* (Winn *et al*., 2011), and those within 0.5 Å distance from any idealized riding position were identified as potential hydrogen atoms. In this manner, we located 376 out of 1067 possible hydrogen atoms corresponding to about 35% of the entire structure. Lowering the threshold to 1.0σ revealed a total of 562 hydrogen atom positions, approximately 53%. At contour levels below 2.0σ, the difference map is nosier, increasing the chance of false positives and making it more challenging to unambiguously identify peaks as hydrogen atoms. Nevertheless, these results are the most complete hydrogen bonding network visualized to date by macromolecular MicroED (Table 1).

**Table 1.** Hydrogen atoms and mean observed hydrogen bond lengths.

| Hydrogen bonds | No. observations | X-H (Å)<sup>a</sup> | X-H<sub>X-ray</sub> (Å)<sup>b</sup> | X-H<sub>Neutron</sub> (Å)<sup>c</sup> |
| --- | --- | --- | --- | --- |
| C<sub>α</sub>-H | 61 | 1.11(13) | 0.98 | 1.10 |
| C<sub>sidechain</sub>-H | 14 | 1.25(18) | 0.98 | 1.10 |
| C<sub>aromatic</sub>-H | 17 | 1.13(12) | 0.93 | 1.08 |
| C-H<sub>2</sub> | 99 | 1.17(15) | 0.97 | 1.09 |
| C-H<sub>3</sub> | 77 | 1.09(17) | 0.96 | 1.06 |
| N-H | 44 | 1.02(16) | 0.86 | 1.01 |
| N-H...O | 38 | 1.05(15) | 0.86 | 1.01 |
| N-H<sub>2</sub> | 13 | 1.08(21) | 0.89 | 1.03 |
| N-H<sub>3</sub> | 3 | 1.13(11) | 0.86 | 1.01 |
| O-H | 10 | 1.13(18) | 0.82 | 0.98 |

<sup>a</sup> Mean observed hydrogen bond lengths measured for hydrogen atoms difference peaks at $\geq 2.0\sigma$ , standard deviations are listed in parenthesis. Values for individual hydrogen bond distances are listed in Supplementary Tables 2–10
<sup>b</sup> Idealized hydrogen bond lengths between electron cloud centroids used in X-ray diffraction (Gruene *et al.*, 2014)
<sup>c</sup> Idealized inter-nuclei hydrogen bond lengths used in neutron diffraction (Gruene *et al.*, 2014)

## Visualizing hydrogens and hydrogen bonding networks

Overall, the protein main chain is expected to be more rigid than the side chains; we consequently expect more hydrogen atoms to be found in the backbone than in the protein side chains. At the 2.0σ threshold, we identified 61 out of 141 possible Cα-H hydrogens and 76 out of 127 peptide N-H hydrogen bonds corresponding to approximately 43 and 60% of the entire structure, respectively (Table 1, Supplementary Tables 2, 3, and 8). The backbone hydrogen atoms are structurally important and can be involved in forming and stabilizing secondary structural elements via non-covalent hydrogen-bonding interactions. For example, the structure of lysozyme has two short antiparallel β-strands and we could identify three strong difference peaks at >3.0σ indicating the positions of those hydrogen atoms involved in hydrogen bonding interactions (Fig. 1a, Supplementary Video 1). The average N-H distance in the β-strands is 1.14(26) Å distance, and the distance between the amide group hydrogen donor and carbonyl acceptor is 2.76(9) Å (Table 2). Interestingly, whereas the Asp52 and Gly54 N-H distances are close to the idealized positions, the difference peak for the Asn44 N-H is located at an almost equal distance shared between the donor and Asp52 carbonyl acceptor (Fig. 1a, Table 2). The structure of lysozyme is further composed of several short helices, and we could identify a total of 15 hydrogen bonding interactions in the three standard α-helices (Table 2). For example, in the longest 12-residue α-helix we identified 6 out of 10 possible hydrogen bonds based on strong difference peaks at >2.7σ (Fig. 1b, Supplementary Video 2). The average hydrogen atom peptide N-H distance for the α-helices is 0.97(14) Å with an average distance between donor and acceptor of 2.84(13) Å (Table 2).

**Figure 1.**
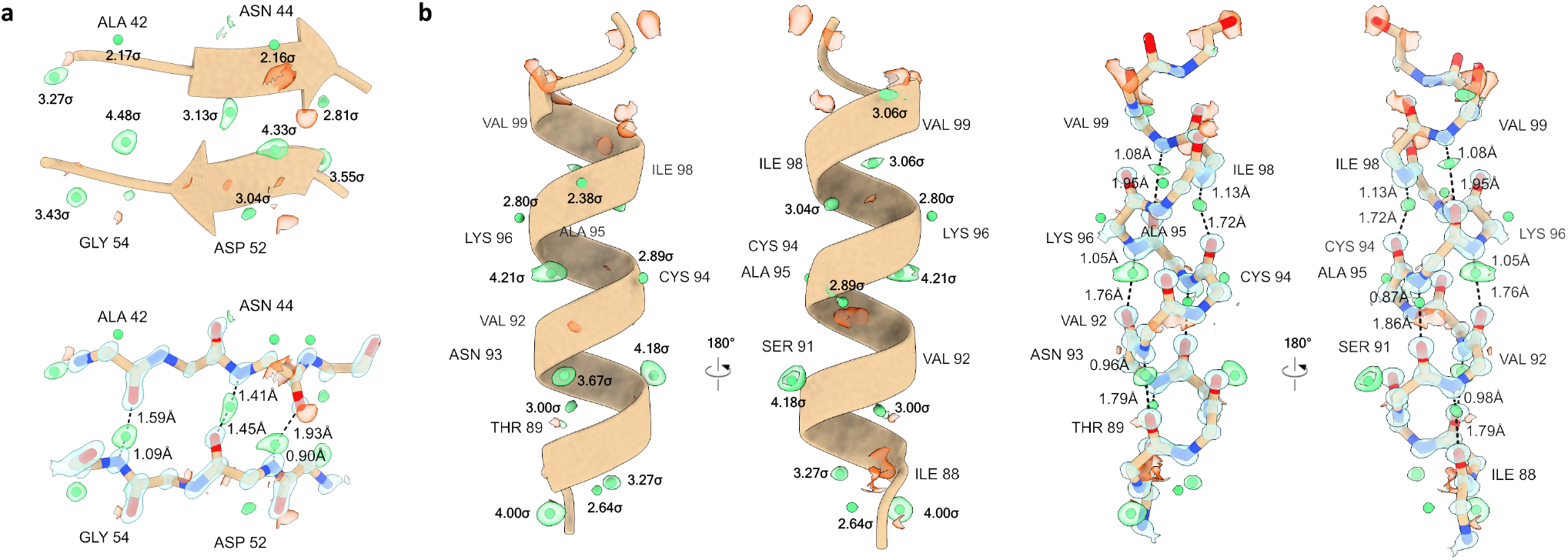
Hydrogen atoms and bonding interactions in secondary structure elements. Difference peaks for individual hydrogen atoms are displayed as green spheres with their σ values shown for (a) two short anti-parallel β-strands (residues 42-45 and 51-54, respectively), and (b) an α-helix (residues 88-101). Hydrogen atoms were assigned from a hydrogen-only omit map for peaks at ≥2.0σ that are within 0.5 Å distance from their idealized position. Hydrogen bonding interactions are indicated by dashed black lines and their respective bond distances and angles are listed in Table 2. Electrostatic potential 2mFo-DFc maps are contoured at 4.0σ (blue) and mFo-DFc difference maps are shown at 2.5σ (green and red for positive and negative, respectively). Carbon atoms are shown in brown, nitrogen in blue, and oxygen in red.

**Table 2.**
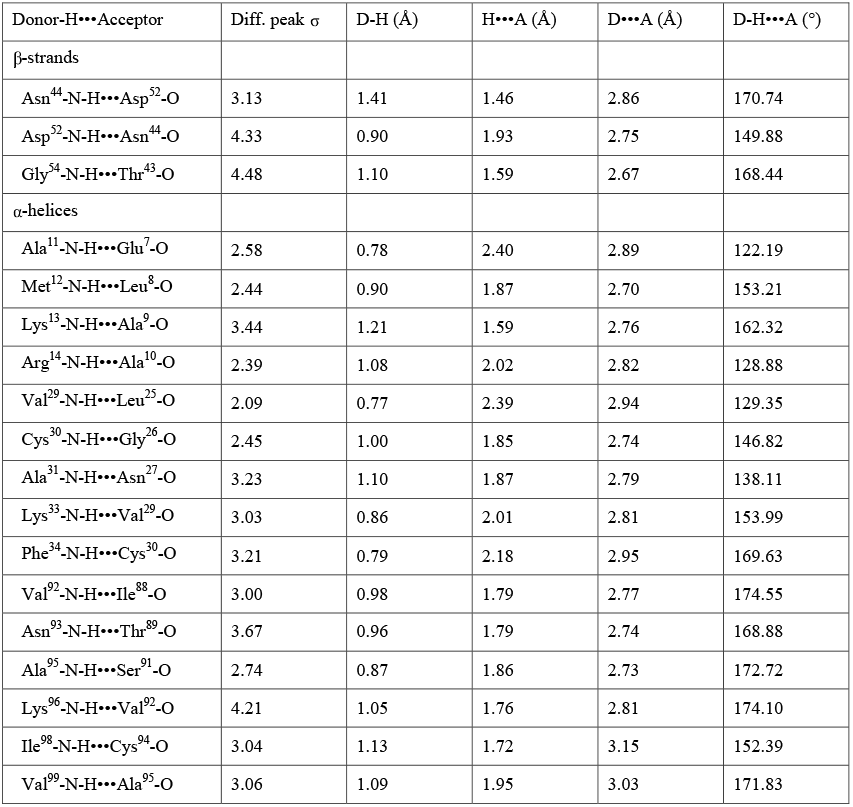
Hydrogen bond distances and angles for secondary structure.

| Donor-H...Acceptor | Diff. peak $\sigma$ | D-H (Å) | H...A (Å) | D...A (Å) | D-H...A (°) |
| --- | --- | --- | --- | --- | --- |
| β-strands | | | | | |
| Asn<sup>44</sup>-N-H...Asp<sup>52</sup>-O | 3.13 | 1.41 | 1.46 | 2.86 | 170.74 |
| Asp<sup>52</sup>-N-H...Asn<sup>44</sup>-O | 4.33 | 0.90 | 1.93 | 2.75 | 149.88 |
| Gly<sup>54</sup>-N-H...Thr<sup>43</sup>-O | 4.48 | 1.10 | 1.59 | 2.67 | 168.44 |
| α-helices | | | | | |
| Ala<sup>11</sup>-N-H...Glu<sup>7</sup>-O | 2.58 | 0.78 | 2.40 | 2.89 | 122.19 |
| Met<sup>12</sup>-N-H...Leu<sup>8</sup>-O | 2.44 | 0.90 | 1.87 | 2.70 | 153.21 |
| Lys<sup>13</sup>-N-H...Ala<sup>9</sup>-O | 3.44 | 1.21 | 1.59 | 2.76 | 162.32 |
| Arg<sup>14</sup>-N-H...Ala<sup>10</sup>-O | 2.39 | 1.08 | 2.02 | 2.82 | 128.88 |
| Val<sup>29</sup>-N-H...Leu<sup>25</sup>-O | 2.09 | 0.77 | 2.39 | 2.94 | 129.35 |
| Cys<sup>30</sup>-N-H...Gly<sup>26</sup>-O | 2.45 | 1.00 | 1.85 | 2.74 | 146.82 |
| Ala<sup>31</sup>-N-H...Asn<sup>27</sup>-O | 3.23 | 1.10 | 1.87 | 2.79 | 138.11 |
| Lys<sup>33</sup>-N-H...Val<sup>29</sup>-O | 3.03 | 0.86 | 2.01 | 2.81 | 153.99 |
| Phe<sup>34</sup>-N-H...Cys<sup>30</sup>-O | 3.21 | 0.79 | 2.18 | 2.95 | 169.63 |
| Val<sup>92</sup>-N-H...Ile<sup>88</sup>-O | 3.00 | 0.98 | 1.79 | 2.77 | 174.55 |
| Asn<sup>93</sup>-N-H...Thr<sup>89</sup>-O | 3.67 | 0.96 | 1.79 | 2.74 | 168.88 |
| Ala<sup>95</sup>-N-H...Ser<sup>91</sup>-O | 2.74 | 0.87 | 1.86 | 2.73 | 172.72 |
| Lys<sup>96</sup>-N-H...Val<sup>92</sup>-O | 4.21 | 1.05 | 1.76 | 2.81 | 174.10 |
| Ile<sup>98</sup>-N-H...Cys<sup>94</sup>-O | 3.04 | 1.13 | 1.72 | 3.15 | 152.39 |
| Val<sup>99</sup>-N-H...Ala<sup>95</sup>-O | 3.06 | 1.09 | 1.95 | 3.03 | 171.83 |

Higher flexibility and alternate conformations can affect localizing hydrogen atoms in the side chains. Nevertheless, we could successfully localize side-chain hydrogen atoms in the data and identify several hydrogen-bonding interactions between side-chain atoms (Fig. 2, Supplementary Table 2). For example, a difference peak at 2.4σ can be resolved between His15-NE2 and Thr89-OG1 indicating a possible shared hydrogen bond between both side chains (Fig. 2a). As expected at pH 4.5, the data shows the solvent-exposed histidine to be protonated at ND1, although the hydrogen distance and angle are different from idealized geometry (Fig. 2a). Another example of hydrogen bonding interactions is illustrated for Tyr53-OH acting as a hydrogen donor to Asp66-OD1 with a strong difference peak at 3.4σ (Fig. 2b, Supplementary Table 2).

**Figure 2.**
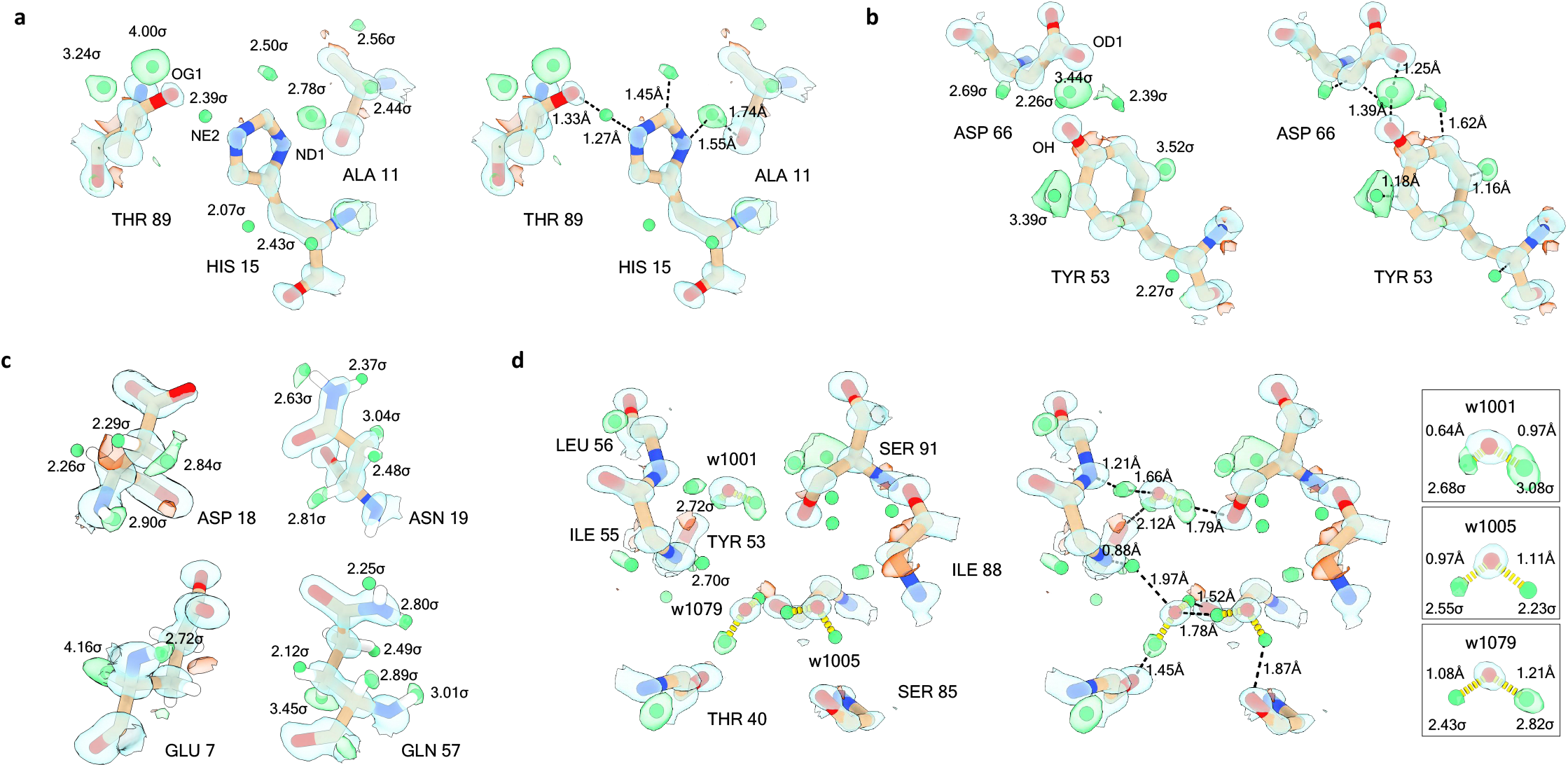
Hydrogen atoms and hydrogen-bond networks. Hydrogen atoms (green difference peaks) are shown with their σ values for the side chains of different residues and for several water molecules. Hydrogen atoms were assigned from a hydrogen-only omit map for peaks at ≥2.0σ and within 0.5Å from their idealized positions. (a) Strong difference peaks indicate hydrogen atom positions for His15, as well as a possible hydrogen bond interaction between His15-NE2 and a neighboring Thr89-OG1. The histidine residue appears to be protonated at ND1 which is consistent with pH 4.5 of the crystallization condition. (b) Hydrogen atoms are indicated by difference peaks for two residues, as well as a potential hydrogen bonding interaction between Tyr53-OH and Asp66-OD1. (c) Acidic side-chains showing well resolved atoms. Strong difference peaks for side chain hydrogen atoms can be observed in asparagine and glutamine residues. (d) Illustration of a hydrogen bonding network involving water molecules and several protein residues. The inset shows the hydrogen bond distances for the water molecules. Electrostatic potential 2mFo-DFc maps are contoured at (a) 2.5σ (blue) and (b-d) at 3.0σ (blue), mFo-DFc difference maps are shown at 2.3σ (green and red for positive and negative, respectively). Carbon atoms are shown in brown, nitrogen in blue, and oxygen in red.

In single-particle cryo-EM, it was previously observed that acidic side chains were poorly resolved at moderate to low resolution owing to radiation damage and due to the rapid fall off of the electron scattering factors for negatively charged atoms at lower scattering angles (Yonekura *et al*., 2015, 2018; Maki-Yonekura *et al*., 2021). In the MicroED data, the acidic aspartate and glutamate residues and their negatively charged side-chain carboxyl groups are generally well resolved (Fig. 2c). Additionally, clear difference peaks at >2.3σ were identified in the data for the amide side-chain nitrogen for asparagine and glutamine residues, making it possible to clearly distinguish between the nitrogen and oxygen atoms of the side-chain amide group (Fig. 2c).

Difference peaks were also identified for several water molecules that are involved in hydrogen bonding interactions with the protein backbone and side chains (Fig. 2d, Supplementary Video 3). Such hydrogen bonding networks can act as long-range proton transfer wires. For example, a water molecule is coordinated with the adjacent Ser91, Leu56, and Tyr53 residues and shows two strong difference peaks at ≥2.7σ (Fig. 2d). Two additional water molecules show hydrogen atom peaks at ≥2.2σ and are involved in hydrogen bonding interactions with each other and residues of the neighboring protein backbone (Fig. 2d). The O-H hydrogen bond lengths and angles of the water molecules are reasonably close to ideal values, except for one O-H distance for w1001 which is significantly shorter at 0.64 Å. The hydrogen bond distance between the w1001-O proton donor and the Tyr53-O proton acceptor is however close to ideal values at 2.75 Å.

## Hydrogen bond distances

The sheer numbers of hydrogens visualized in this study allow us to measure and report hydrogen bond distances in a way previously not possible in cryoEM (Supplementary Tables 2–10; Figure 3). Electrons are scattered by the potential field generated from electron clouds *and* the nuclei; similar to neutron diffraction. The peaks in an electrostatic potential map are therefore expected to reflect the inter-nuclei distances more than distances between centers of mass of electron clouds as observed in X-ray diffraction. We refined the structure using the default riding hydrogen model based on hydrogen distances between the electron cloud centroids using restraints derived from X-ray scattering. We analyzed the identified hydrogen atom difference peaks in the data at ≥2.0σ and calculated the average distance for each of the hydrogen bond types (Table 1, Figure 3, Supplementary Tables 2–10). The number of observations for some bond types is insufficient for a rigorous statistical analysis. We do however find an average Cα-H distance for the main chain of 1.11(13) Å for 61 hydrogen bonds, compared to idealized values of 0.98 and 1.10 Å for X-ray and neutron diffraction, respectively (Table 1, Figure 3, Supplementary Table 3). The average distance for all N-H bonds is 1.03(16) Å for 83 observations, compared to idealized values of 0.98 and 1.10 Å for X-ray and neutron diffraction, respectively (Supplementary Tables 2 and 8). Interestingly, the distances for the amide N-H bonds that are involved in hydrogen bonding interactions with neighboring residues are slightly longer compared to those that are not involved in such electrostatic interactions (Table 1, Figure 3).

**Figure 3.**
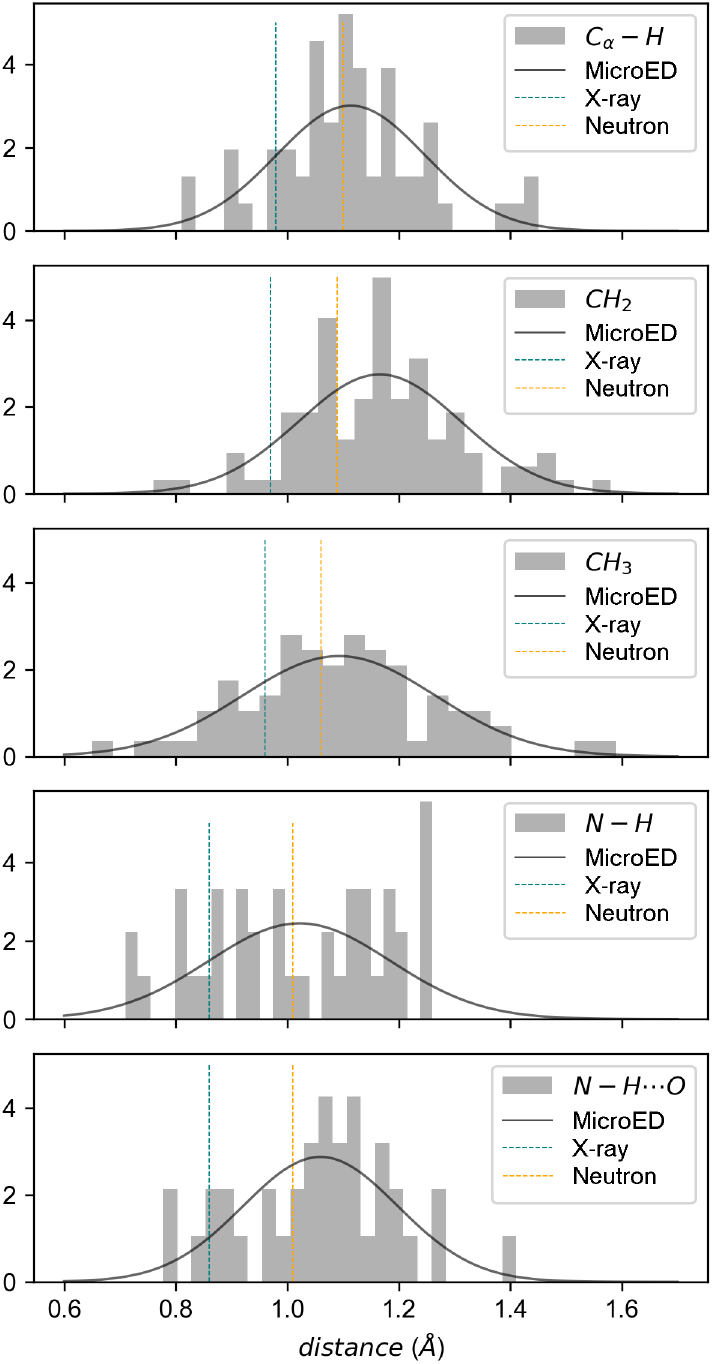
Hydrogen bond distances for macromolecular MicroED data. Hydrogen bond distances in Å are shown as histogram plots with a normal distribution fitted to the data. Idealized hydrogen bond lengths between electron cloud centroids used in X-ray diffraction are indicated by a teal dotted line, idealized inter-nuclei hydrogen bond lengths used in neutron diffraction are indicated by the orange dotted line (see also Table 1, Supplementary Tables 2, 3, 6–8) (Gruene *et al*., 2014).

These results suggest an elongation of the hydrogen bond lengths compared to the electron cloud centroid distances assumed in the riding hydrogen mode, although the number of observations for each type is rather limited and the standard deviations from the mean value are quite large (Table 1, Figure 3). Nevertheless, we find an overall trend that the Cα-H and N-H bond lengths are closer inter-nuclei distances (Gruene *et al*., 2014; Williams *et al*., 2018). This observation agrees with previous electron diffraction and imaging experiments that show an apparent elongation of the hydrogen bond lengths compared to X-ray diffraction (Clabbers *et al*., 2019; Takaba *et al*., 2021; Nakane *et al*., 2020; Maki-Yonekura *et al*., 2021). Refinement of structural models derived from electron scattering would therefore benefit from more appropriate restraints specific for electrons, including a more accurate riding hydrogen model, as well as taking the electrostatic potential of the crystal into account (Yonekura *et al*., 2015, 2018; Murshudov *et al*., 2011; Yamashita *et al*., 2021; Gruene *et al*., 2014; Williams *et al*., 2018).

## Conclusions

The results demonstrate that hydrogen atom positions can be accurately identified in macromolecular MicroED data. As with X-ray crystallography, this will typically require atomic resolution data or better (Walsh *et al*., 1998; Howard *et al*., 2004; Wang *et al*., 2007; Eriksson *et al*., 2013; Ogata *et al*., 2015). In comparison, the structure of triclinic lysozyme was determined previously using X-ray diffraction at 120 K and room temperature to 0.93 and 0.95 Å resolution, respectively (Walsh *et al*., 1998). The single-crystal low-temperature structure is of high quality and generally has more clearly visible hydrogen atoms than the room temperature model merged from three crystal datasets. Difference maps contoured at 1.9σ visualize hydrogen atoms in residues within the better-defined regions of the structure, and at 1.8σ contour level, 77 out of 127 peptide N-H atoms (61%) are identified (Walsh *et al*., 1998). The number of hydrogen atoms localized in the low-temperature structure is similar to the MicroED structure at comparable resolution, even though the intensity and model statistics are worse (Supplementary Table 1, Supplementary Fig. 1) (Martynowycz, Clabbers, Hattne *et al*., 2021). The higher level of inaccuracy in the MicroED data can in part be attributed to non-isomorphism from merging of 16 crystal datasets and lower completeness in the highest resolution shells (Supplementary Fig. 1). Additional factors that contribute to the errors are inelastic scattering and inaccurate modeling of the electron form factors and the electrostatic potential in structure refinement (Yonekura *et al*., 2015, 2018; Murshudov *et al*., 2011; Yamashita *et al*., 2021). Compared to X-ray diffraction, electrons are expected to provide better contrast for identifying hydrogen atoms at a similar resolution as the scattering factors fall off less steeply with decreasing atomic number. The lighter hydrogen atoms are therefore expected to be better resolved next to the heavier atoms, which might explain why we can identify many hydrogen atoms even though the MicroED data appear noisier. This is further supported by a comparison between apoferritin models from X-ray crystallography and single-particle cryo-EM showed that hydrogen atoms were visibly more clearly in the latter (Yamashita *et al*., 2021). More recently, a significantly higher resolution structure of triclinic lysozyme was solved *ab initio* at 0.65 Å by X-ray diffraction (Wang *et al*., 2007). At this resolution, approximately 31% of all hydrogen atoms in main and side chains could be identified at 3.0σ or higher. We would anticipate major improvements in hydrogen atom localization in MicroED data upon further increasing the resolution.

Previously, hydrogen atoms were successfully identified in protein complexes by single-particle cryo-EM. In comparison to the results presented here, these studies reported that about 70% of the expected number of hydrogen atoms could be identified above a threshold level of 2.0σ using hydrogen-only omit maps from atomic resolution reconstructions of apoferritin (Yamashita *et al*., 2021; Maki-Yonekura *et al*., 2021). Remarkably, about 17% of possible hydrogen atoms could be identified from data as low as 1.84 Å resolution (Yamashita *et al*., 2021). In imaging, the phase information is retained and during reconstruction, images are filtered to remove noise and to select a specific conformational state. The resolution is therefore a local feature of the map whereas the *B*-factor is a global parameter applied in map sharpening or blurring. This is unlike a crystallographic map, where the resolution is a global feature of the entire dataset and structural flexibility or disorder is modeled locally using alternate conformation and per atom refined *B*-factors. In crystallography, resolving detailed features such as hydrogen atoms is affected by local disorder. In the MicroED structure, twelve residues are modeled with alternate side-chain conformations at low occupancy, making it more challenging to identify hydrogen at these positions. The mean *B*-factor over all atoms in the model is 11.98 Å^2^, and the majority of outliers are on the outside of the protein facing the solvent. Lower completeness in the higher resolution shells and non-isomorphism from merging data of multiple crystals could both contribute to increased *B*-factors. Especially the last two C-terminal residues have high temperature factors, these were also poorly resolved in the high-resolution X-ray diffraction structure (Walsh *et al*., 1998). Indeed, most hydrogens can be identified within the more stable core of the protein relative to the residues on the outside facing the solvent having higher flexibility and *B*-factors.

In all, 377/1067 (35%) hydrogen atoms could be located at ≥2.0σ and we illustrate several examples of well resolved hydrogen atom positions and hydrogen bonding interactions between protein residues and solvent molecules. This is the most complete hydrogen network map for macromolecular MicroED data to date, and these results provide a glimpse of the information that can be obtained by electron scattering, opening up new avenues for further experiments investigating hydrogen bonding networks in protein structures. At the current stage, the difference map becomes increasingly noisy at contour levels below 2.0σ, making it more challenging to unambiguously identify hydrogen peaks. Future efforts that can enhance the localization of hydrogen atoms should be focused on improving data accuracy and resolution even further. Energy filtration can improve data quality by discarding inelastically scattered electrons, improving the detection of weak peaks at high resolution and at the lower scattering angles that are shaded by the direct beam (Yonekura *et al*., 2015, 2019). It would also mean the exposure could be lowered even further without losing the weak signal from high-resolution reflections to the noise of the background. Energy filtering does not exclude multiple elastic scattering which may affect the measured kinematic intensities (Fujiwara, 1959; Cowley, 1995). For any typical hydrated protein crystal, these effects are suggested to be far less detrimental to data quality compared to inelastic scattering (Latychevskaia & Abrahams, 2019; Martynowycz, Clabbers, Unge *et al*., 2021). Dynamical structure refinement can enhance the localization of hydrogen atoms in small molecule structures (Palatinus *et al*., 2017), but its implementation is computationally expensive and has yet to be extended to macromolecules that include bulk solvent that cannot be modeled. In recent experiments, recording MicroED data using a direct electron detector in electron counting mode significantly improved data quality, and we expect further benefits from faster readout and better electron-counting algorithms using electron-event representation (Guo *et al*., 2020; Nakane *et al*., 2020).

## Methods

### Crystallization and sample preparation

Crystalline lamellae of triclinic lysozyme were prepared as described previously (Martynowycz, Clabbers, Hattne *et al*., 2021). Briefly, crystals of hen egg-white lysozyme (*Gallus gallus*) were grown by dissolving 10 mg/ml protein in a solution of 0.2 M sodium nitrate and 50 mM sodium acetate at pH 4.5. After incubation overnight at 4 °C an opaque suspension was observed. After further incubation for one week at room temperature a crystalline slurry appeared containing microcrystals. Samples were prepared by depositing 3 μl of the crystalline slurry onto a glow-discharged EM grid (Quantifoil, Cu 200 mesh, R2/2 holey carbon). Excess liquid was blotted away and the sample was vitrified using a Leica GP2 vitrification robot. Grids were transferred to an Aquilos dual-beam FIB/SEM (Thermo Fisher) and crystals were milled to lamellae with an optimal thickness of approximately 300 nm as described previously (Martynowycz, Clabbers, Hattne *et al*., 2021; Martynowycz, Clabbers, Unge *et al*., 2021).

### Data collection and processing

Electron-counted MicroED data were collected on a Titan Krios 3Gi TEM (Thermo Fisher) operated at 300 kV as described previously (Martynowycz, Clabbers, Hattne *et al*., 2021). Briefly, the TEM was set up for low exposure data collection using a 50 μm C2 aperture, spot size 11, and a beam diameter of 25 μm. A 100 μm SA aperture was used, corresponding to an area of 2 μm diameter on the specimen. Crystal lamellae were continuously rotated over a range of 84° at a rotation speed of 0.2°/s over 420s with a total exposure of approximately 0.64 e^-^.Å^-2^ per dataset. Data were recorded on a Falcon 4 direct electron detector (Thermo Fisher) in electron counting mode operating at an internal frame rate of 250 Hz. Data of 16 crystal lamellae were integrated, scaled and merged using *XDS* (Kabsch, 2010) and *AIMLESS* (Evans & Murshudov, 2013). The structure was phased *ab initio* by placing a three-residue idealized α-helix fragment using *PHASER* (McCoy *et al*., 2007) followed by density modification in *ACORN* (Foadi *et al*., 2000). The entire structure was built automatically using *BUCCANEER* (Cowtan, 2006) and refined in *REFMAC5* (Murshudov *et al*., 2011) using electron scattering factors.

### Identification of hydrogen atoms

The structure was manually inspected and remodeled using *Coot* (Emsley *et al*., 2010), and re-refined with *REFMAC5* (Murshudov *et al*., 2011) using electron scattering factors. Hydrogen atoms were added in idealized riding positions. A hydrogen-only omit was calculated from the final structural model by *REFMAC5* (Murshudov *et al*., 2011). Peaks in the *mF_o_* – *DF_C_* difference map at a threshold above 2.0σ were identified and listed using *PEAKMAX* in the CCP4 software package (Winn *et al*., 2011). Difference peaks that fell within 0.5 Å of the idealized distance for the known positions were assigned as hydrogen atoms.

### Figure preparation

Figures were prepared using ChimeraX and assembled in powerpoint and photoshop.

## Data availability

Coordinates and structure factors have been deposited to the PDB.

## Acknowledgements

This study was supported by the National Institutes of Health P41GM136508. The Gonen laboratory is supported by funds from the Howard Hughes Medical Institute.

---

**Supplementary Figure 1.**
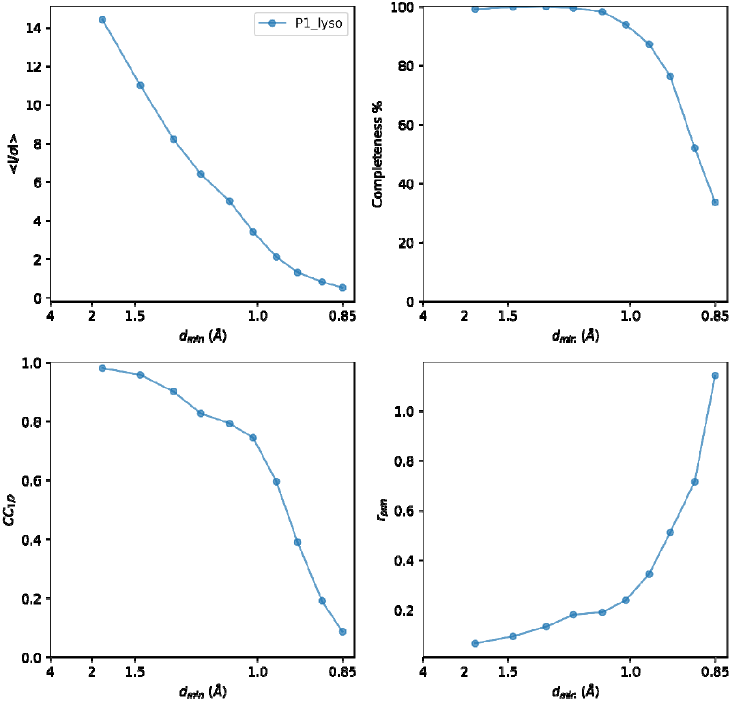
Merging statistics for triclinic lysozyme. Crystallographic quality indicators and data completeness plotted as function of resolution for triclinic lysozyme at 0.87 Å resolution (see Supplementary Table 1).

**Supplementary Table 1.** MicroED Data collection and refinement statistics.

| **Data collection** * |  |
| --- | --- |
| Wavelength | 0.0197 |
| No. of crystals | 16 |
| Space group | *P*1 |
| Cell dimensions |  |
| *a*, *b*, *c* (Å) | 26.42, 30.72, 33.01 |
| $\alpha$, $\beta$, $\gamma$ (°) | 88.32, 109.10, 112.08 |
| Resolution (Å) | 16.05-0.87 (0.90-0.87)** |
| Observed reflections | 569407 (5797) |
| Unique reflections | 64986 (2783) |
| Multiplicity | 8.8 (2.1) |
| Completeness (%) | 87.55 (37.64) |
| R<sub>merge</sub> | 0.236 (1.035) |
| R<sub>meas</sub> | 0.248 (1.409) |
| R<sub>pim</sub> | 0.073 (0.945) |
| Mean I/σ(I) | 6.23 (0.66) |
| CC<sub>1/2</sub> | 0.990 (0.147) |
| **Refinement** |  |
| No. of reflections | 64974 |
| No. of reflections used for R<sub>free</sub> | 3168 |
| R<sub>work</sub> / R<sub>free</sub> | 0.197 / 0.220 |
| No. of atoms | 1216 |
| Proteins | 1081 |
| Ligand | 16 |
| Water | 119 |
| R.m.s. deviations |  |
| Bond lengths (Å) | 0.034 |
| Bond angles (°) | 2.447 |
| Mean *B*-factor (Å<sup>2</sup>) | 11.98 |
| Ramachandran |  |
| Favored (%) | 96.19 |
| Allowed (%) | 3.81 |
| Outliers (%) | 0.00 |
| Rotamer outliers (%) | 2.33 |

\*Data from Martynowycz, Clabbers, Hattne *et al.*, 2021.
\*\*Values in parentheses are for highest-resolution shell.

**Supplementary Table 2.** Bond distances and angles for hydrogen bonding interactions.

| Donor-H...Acceptor | Diff. peak $\sigma$ | D-H (Å) | H...A (Å) | D...A (Å) | D-H...A (°) |
| --- | --- | --- | --- | --- | --- |
| Phe<sup>3</sup>-N-H...Phe<sup>38</sup>-O | 2.50 | 1.18 | 1.67 | 2.81 | 160.72 |
| Ala<sup>11</sup>-N-H...Glu<sup>7</sup>-O | 2.58 | 0.78 | 2.40 | 2.89 | 122.19 |
| Met<sup>12</sup>-N-H...Leu<sup>8</sup>-O | 2.44 | 0.90 | 1.87 | 2.70 | 153.21 |
| Lys<sup>13</sup>-N-H...Ala<sup>9</sup>-O | 3.44 | 1.21 | 1.59 | 2.76 | 162.32 |
| Arg<sup>14</sup>-N-H...Ala<sup>10</sup>-O | 2.39 | 1.08 | 2.02 | 2.82 | 128.88 |
| His<sup>15</sup>-NE2-H...Thr<sup>89</sup>-OG1 | 2.39 | 1.27 | 1.33 | 2.59 | 162.48 |
| Leu<sup>17</sup>-N-H...Met<sup>12</sup>-O | 2.91 | 1.11 | 1.92 | 2.99 | 162.18 |
| Tyr<sup>20</sup>-N-H...Leu<sup>17</sup>-O | 2.78 | 1.06 | 1.98 | 2.92 | 146.41 |
| Val<sup>29</sup>-N-H...Leu<sup>25</sup>-O | 2.09 | 0.77 | 2.39 | 2.94 | 129.35 |
| Cys<sup>30</sup>-N-H...Gly<sup>26</sup>-O | 2.45 | 1.00 | 1.85 | 2.74 | 146.82 |
| Ala<sup>31</sup>-N-H...Asn<sup>27</sup>-O | 3.23 | 1.10 | 1.87 | 2.79 | 138.11 |
| Lys<sup>33</sup>-N-H...Val<sup>29</sup>-O | 3.03 | 0.86 | 2.01 | 2.81 | 153.99 |
| Phe<sup>34</sup>-N-H...Cys<sup>30</sup>-O | 3.21 | 0.79 | 2.18 | 2.95 | 169.63 |
| Thr<sup>40</sup>-N-H...Lys<sup>1</sup>-O | 3.03 | 1.08 | 1.77 | 2.75 | 147.63 |
| Ala<sup>42</sup>-N-H...Asn<sup>39</sup>-O | 3.27 | 1.20 | 1.77 | 2.89 | 154.32 |
| Asn<sup>44</sup>-N-H...Asp<sup>52</sup>-O | 3.12 | 1.41 | 1.46 | 2.86 | 170.74 |
| Asn<sup>46</sup>-N-H...Ser<sup>50</sup>-O | 2.56 | 1.21 | 1.63 | 2.73 | 148.76 |
| Asp<sup>52</sup>-N-H...Asn<sup>44</sup>-O | 4.33 | 0.90 | 1.93 | 2.75 | 149.88 |
| Tyr<sup>53</sup>-O-H...Asp<sup>66</sup>-OD2 | 3.44 | 1.39 | 1.25 | 2.59 | 157.86 |
| Gly<sup>54</sup>-N-H...Thr<sup>43</sup>-O | 4.48 | 1.10 | 1.59 | 2.67 | 168.44 |
| Gln<sup>57</sup>-N-H...Gly<sup>54</sup>-O | 3.01 | 1.11 | 1.83 | 2.84 | 150.17 |
| Asn<sup>65</sup>-N-H...Leu<sup>78</sup>-O | 2.74 | 0.91 | 1.87 | 2.77 | 159.36 |
| Gly<sup>67</sup>-N-H...Asn<sup>65</sup>-O | 3.90 | 1.13 | 1.85 | 2.90 | 152.42 |
| Arg<sup>73</sup>-N-H...Arg<sup>61</sup>-O | 3.77 | 1.17 | 1.74 | 2.80 | 147.11 |
| Leu<sup>75</sup>-N-H...Trp<sup>62</sup>-O | 2.86 | 0.84 | 1.93 | 2.75 | 161.86 |
| Ser<sup>81</sup>-N-H...NO3<sup>201</sup>-O | 2.92 | 1.07 | 1.8 | 2.76 | 146.94 |
| Ala<sup>82</sup>-N-H...Pro<sup>70</sup>-O | 2.16 | 1.18 | 1.71 | 2.79 | 148.60 |
| Leu<sup>83</sup>-N-H...Cys<sup>80</sup>-O | 2.32 | 1.04 | 1.77 | 2.73 | 150.33 |
| Leu<sup>84</sup>-N-H...Ser<sup>81</sup>-O | 2.99 | 1.03 | 1.92 | 2.93 | 164.49 |
| Val<sup>92</sup>-N-H...Ile<sup>88</sup>-O | 3.00 | 0.98 | 1.79 | 2.77 | 174.55 |
| Asn<sup>93</sup>-N-H...Thr<sup>89</sup>-O | 3.67 | 0.96 | 1.79 | 2.74 | 168.89 |
| Ala<sup>95</sup>-N-H...Ser<sup>91</sup>-O | 2.74 | 0.87 | 1.86 | 2.73 | 172.72 |
| Lys<sup>96</sup>-N-H•••Val<sup>92</sup>-O | 4.21 | 1.05 | 1.76 | 2.81 | 174.10 |
| Ile<sup>98</sup>-N-H•••Cys<sup>94</sup>-O | 3.04 | 1.13 | 1.72 | 3.15 | 152.39 |
| Val<sup>99</sup>-N-H•••Ala<sup>95</sup>-O | 3.06 | 1.09 | 1.95 | 3.03 | 171.83 |
| Trp<sup>108</sup>-N-H•••Met<sup>105</sup>-O | 3.10 | 1.26 | 1.64 | 2.82 | 152.36 |
| Trp<sup>111</sup>-NE1-H•••Asn<sup>27</sup>-OD1 | 3.39 | 1.02 | 1.79 | 2.71 | 169.48 |
| Arg<sup>114</sup>-N-H•••Arg<sup>110</sup>-O | 2.27 | 1.11 | 1.98 | 2.76 | 124.52 |
| Trp<sup>123</sup>-N-H•••Val<sup>120</sup>-O | 2.45 | 1.04 | 1.83 | 2.84 | 161.33 |

**Supplementary Table 3.** Hydrogen bond distances for Cα-H.

| Residue | Name | Atom | Diff. peak $\sigma$ | X-H (Å) |
| --- | --- | --- | --- | --- |
| 2 | Val | CA | 3.42 | 1.15 |
| 3 | Phe | CA | 3.46 | 1.26 |
| 4 | Gly | CA | 3.18 | 1.14 |
| 4 | Gly | CA | 2.39 | 0.98 |
| 5 | Arg | CA | 2.59 | 1.27 |
| 7 | Glu | CA | 4.16 | 1.06 |
| 9 | Ala | CA | 3.79 | 1.08 |
| 10 | Ala | CA | 2.74 | 1.00 |
| 11 | Ala | CA | 2.78 | 0.81 |
| 12 | Met | CA | 2.74 | 1.11 |
| 13 | Lys | CA | 2.01 | 1.20 |
| 16 | Gly | CA | 2.60 | 0.89 |
| 18 | Asp | CA | 2.26 | 1.45 |
| 19 | Asn | CA | 2.81 | 1.43 |
| 20 | Tyr | CA | 3.17 | 1.10 |
| 21 | Arg | CA | 3.12 | 1.29 |
| 22 | Gly | CA | 2.77 | 1.18 |
| 24 | Ser | CA | 3.96 | 1.10 |
| 25 | Leu | CA | 3.06 | 1.19 |
| 27 | Asn | CA | 3.22 | 1.16 |
| 28 | Trp | CA | 3.34 | 1.10 |
| 29 | Val | CA | 2.42 | 1.10 |
| 32 | Ala | CA | 2.39 | 1.05 |
| 33 | Lys | CA | 2.71 | 1.05 |
| 38 | Phe | CA | 2.41 | 1.18 |
| 39 | Asn | CA | 2.96 | 0.99 |
| 42 | Ala | CA | 2.17 | 0.83 |
| 44 | Asn | CA | 2.16 | 0.90 |
| 45 | Arg | CA | 2.81 | 0.98 |
| 51 | Thr | CA | 3.55 | 0.90 |
| 52 | Asp | CA | 3.04 | 1.20 |
| 53 | Tyr | CA | 2.27 | 1.14 |
| 54 | Gly | CA | 3.43 | 1.17 |
| 55 | Ile | CA | 2.63 | 1.12 |
| 56 | Ile | CA | 2.85 | 1.25 |
| 57 | Asn | CA | 2.89 | 1.23 |
| 58 | Ile | CA | 3.49 | 0.97 |
| 59 | Asn | CA | 3.51 | 1.06 |
| 62 | Trp | CA | 2.78 | 1.18 |
| 63 | Trp | CA | 2.29 | 1.25 |
| 69 | Thr | CA | 2.61 | 1.00 |
| 70 | Pro | CA | 2.91 | 1.06 |
| 74 | Asn | CA | 3.22 | 1.03 |
| 75 | Leu | CA | 2.65 | 1.13 |
| 76 | Cys | CA | 2.76 | 1.08 |
| 77 | Asn | CA | 2.87 | 1.13 |
| 82 | Ala | CA | 3.27 | 1.09 |
| 88 | Ile | CA | 2.64 | 1.41 |
| 91 | Ser | CA | 4.18 | 1.40 |
| 94 | Cys | CA | 2.89 | 1.02 |
| 96 | Lys | CA | 2.80 | 1.05 |
| 97 | Lys | CA | 2.38 | 1.24 |
| 99 | Val | CA | 3.06 | 1.15 |
| 103 | Asn | CA | 2.14 | 1.17 |
| 108 | Trp | CA | 4.03 | 0.94 |
| 110 | Ala | CA | 3.41 | 1.06 |
| 111 | Trp | CA | 2.44 | 1.10 |
| 116 | Lys | CA | 2.93 | 1.11 |
| 117 | Gly | CA | 4.02 | 1.11 |
| 119 | Asp | CA | 3.19 | 1.07 |
| 123 | Trp | CA | 2.60 | 1.14 |

**Supplementary Table 4.** Hydrogen bond distances for side chain C-H.

| Residue | Name | Atom | Diff. peak $\sigma$ | X-H (Å) |
| --- | --- | --- | --- | --- |
| 8 | Leu | CG | 2.50 | 0.94 |
| 15 | His | CE1 | 2.50 | 1.45 |
| 40 | Thr | CB | 2.64 | 1.51 |
| 56 | Leu | CG | 3.74 | 1.22 |
| 58 | Ile | CB | 3.35 | 1.18 |
| 63 | Trp | CD1 | 3.06 | 1.39 |
| 69 | Thr | CB | 3.08 | 0.99 |
| 84 | Leu | CG | 2.88 | 1.29 |
| 89 | Thr | CB | 3.24 | 1.25 |
| 98 | Ile | CB | 2.19 | 1.06 |
| 108 | Trp | CD1 | 2.84 | 1.25 |
| 109 | Val | CB | 2.36 | 1.51 |
| 118 | Thr | CB | 3.39 | 1.29 |
| 123 | Trp | CD1 | 2.44 | 1.19 |

**Supplementary Table 5.** Hydrogen bond distances for aromatic C-H.

| Residue | Name | Atom | Diff. peak $\sigma$ | X-H (Å) |
| --- | --- | --- | --- | --- |
| 3 | Phe | CE2 | 3.12 | 1.05 |
| 20 | Tyr | CE1 | 3.09 | 0.99 |
| 23 | Tyr | CD2 | 4.18 | 1.01 |
| 23 | Tyr | CD1 | 2.81 | 1.11 |
| 23 | Tyr | CE1 | 2.15 | 1.32 |
| 38 | Phe | CZ | 3.97 | 1.13 |
| 38 | Phe | CE2 | 2.56 | 1.32 |
| 38 | Phe | CD1 | 2.11 | 1.03 |
| 53 | Tyr | CD1 | 3.52 | 1.18 |
| 53 | Tyr | CE2 | 3.39 | 1.16 |
| 63 | Trp | CZ3 | 3.51 | 1.00 |
| 63 | Trp | CE3 | 3.19 | 1.22 |
| 108 | Trp | CE3 | 3.86 | 1.07 |
| 108 | Trp | CH2 | 3.47 | 1.01 |
| 108 | Trp | CZ3 | 2.42 | 1.31 |
| 111 | Trp | CH2 | 3.52 | 1.24 |
| 111 | Trp | CZ2 | 2.30 | 1.02 |

**Supplementary Table 6.** Hydrogen bond distances for CH_2_.

| Residue | Name | Atom | Diff. peak $\sigma$ | X-H (Å) |
| --- | --- | --- | --- | --- |
| 1 | Lys | CE | 3.20 | 1.41 |
| 1 | Lys | CE | 2.30 | 1.17 |
| 1 | Lys | CG | 3.15 | 1.05 |
| 1 | Lys | CB | 3.12 | 1.16 |
| 3 | Phe | CB | 3.37 | 0.98 |
| 3 | Phe | CB | 2.43 | 1.14 |
| 5 | Arg | CB | 3.12 | 1.23 |
| 5 | Arg | CB | 2.81 | 1.46 |
| 5 | Arg | CG | 2.20 | 1.14 |
| 6 | Cys | CB | 2.37 | 1.04 |
| 7 | Glu | CB | 2.42 | 1.58 |
| 8 | Leu | CB | 2.45 | 1.03 |
| 8 | Leu | CB | 2.43 | 1.10 |
| 12 | Met | CG | 3.12 | 1.18 |
| 14 | Arg | CG | 3.82 | 1.00 |
| 15 | His | CB | 2.07 | 1.16 |
| 18 | Asp | CB | 2.84 | 1.00 |
| 18 | Asp | CB | 2.29 | 1.21 |
| 19 | Asn | CB | 3.04 | 1.01 |
| 19 | Asn | CB | 2.48 | 1.06 |
| 20 | Tyr | CB | 2.43 | 1.06 |
| 20 | Tyr | CB | 2.20 | 1.07 |
| 21 | Arg | CG | 3.36 | 1.17 |
| 21 | Arg | CB | 2.38 | 1.16 |
| 21 | Arg | CB | 2.16 | 1.47 |
| 23 | Trp | CB | 2.80 | 1.14 |
| 25 | Leu | CB | 3.22 | 1.08 |
| 27 | Asn | CB | 2.69 | 0.91 |
| 28 | Trp | CB | 2.19 | 1.06 |
| 33 | Lys | CE | 3.45 | 1.07 |
| 33 | Lys | CG | 3.11 | 1.08 |
| 33 | Lys | CG | 2.27 | 1.13 |
| 34 | Phe | CB | 4.37 | 1.18 |
| 35 | Glu | CG | 3.11 | 1.17 |
| 35 | Glu | CG | 2.53 | 1.34 |
| 36 | Ser | CB | 3.51 | 1.09 |
| 36 | Ser | CB | 2.63 | 1.51 |
| 38 | Phe | CB | 3.14 | 1.06 |
| 41 | Gln | CB | 3.12 | 1.19 |
| 41 | Gln | CB | 2.76 | 1.00 |
| 41 | Gln | CG | 2.48 | 1.10 |
| 48 | Asp | CB | 2.11 | 1.29 |
| 57 | Gln | CB | 3.45 | 1.18 |
| 57 | Gln | CB | 2.49 | 1.19 |
| 58 | Ile | CG1 | 3.79 | 1.23 |
| 58 | Ile | CG1 | 2.79 | 1.12 |
| 59 | Asn | CB | 3.11 | 1.15 |
| 59 | Asn | CB | 2.52 | 1.06 |
| 60 | Ser | CB | 3.67 | 1.30 |
| 61 | Arg | CB | 4.18 | 1.04 |
| 61 | Arg | CB | 2.59 | 0.91 |
| 61 | Arg | CG | 2.40 | 1.47 |
| 62 | Trp | CB | 3.11 | 1.25 |
| 63 | Trp | CB | 2.97 | 1.29 |
| 63 | Trp | CB | 2.34 | 1.29 |
| 64 | Cys | CB | 2.55 | 1.07 |
| 66 | Asp | CB | 2.69 | 1.19 |
| 66 | Asp | CB | 2.26 | 1.43 |
| 68 | Arg | CB | 4.31 | 1.16 |
| 70 | Pro | CB | 2.76 | 1.27 |
| 72 | Ser | CB | 2.04 | 1.14 |
| 73 | Arg | CB | 2.22 | 1.18 |
| 74 | Asn | CB | 2.46 | 1.16 |
| 74 | Asn | CB | 2.14 | 1.27 |
| 75 | Leu | CB | 2.82 | 1.35 |
| 76 | Cys | CB | 3.12 | 0.76 |
| 76 | Cys | CB | 3.07 | 1.21 |
| 77 | Asn | CB | 2.76 | 1.08 |
| 78 | Ile | CG1 | 2.13 | 1.05 |
| 80 | Cys | CB | 3.14 | 0.99 |
| 80 | Cys | CB | 2.60 | 1.31 |
| 83 | Leu | CB | 2.57 | 1.01 |
| 84 | Leu | CB | 2.72 | 1.23 |
| 85 | Ser | CB | 2.37 | 1.28 |
| 86 | Ser | CB | 2.91 | 1.22 |
| 87 | Asp | CB | 3.81 | 1.20 |
| 88 | Ile | CG1 | 2.16 | 1.34 |
| 91 | Ser | CB | 2.86 | 0.95 |
| 91 | Ser | CB | 2.10 | 1.16 |
| 93 | Asn | CB | 3.61 | 1.40 |
| 93 | Asn | CB | 2.59 | 1.15 |
| 94 | Cys | CB | 3.46 | 1.08 |
| 94 | Cys | CB | 2.50 | 1.20 |
| 96 | Lys | CB | 3.05 | 0.90 |
| 97 | Lys | CD | 4.05 | 1.25 |
| 97 | Lys | CG | 3.62 | 1.07 |
| 97 | Lys | CG | 2.88 | 1.26 |
| 97 | Lys | CD2 | 2.97 | 1.30 |
| 97 | Lys | CD2 | 2.65 | 1.18 |
| 97 | Lys | CB | 2.60 | 0.82 |
| 100 | Ser | CB | 3.19 | 1.24 |
| 105 | Met | CB | 2.62 | 1.42 |
| 105 | Met | CG | 2.35 | 1.23 |
| 108 | Trp | CB | 3.41 | 1.17 |
| 114 | Arg | CB | 2.26 | 1.16 |
| 116 | Lys | CE | 3.61 | 1.23 |
| 121 | Gln | CB | 2.77 | 1.04 |
| 121 | Gln | CB | 2.50 | 1.23 |
| 123 | Trp | CB | 2.11 | 1.41 |

**Supplementary Table 7.** Hydrogen bond distances for CH_3_.

| Residue | Name | Atom | Diff. peak $\sigma$ | X-H (Å) |
| --- | --- | --- | --- | --- |
| 2 | Val | CG1 | 3.42 | 0.94 |
| 2 | Val | CG1 | 2.23 | 1.15 |
| 2 | Val | CG2 | 3.04 | 1.31 |
| 2 | Val | CG2 | 2.63 | 1.11 |
| 8 | Leu | CD1 | 3.59 | 1.20 |
| 8 | Leu | CD1 | 3.12 | 0.99 |
| 9 | Ala | CB | 2.93 | 1.00 |
| 10 | Ala | CB | 3.73 | 1.12 |
| 10 | Ala | CB | 2.89 | 1.54 |
| 10 | Ala | CB | 2.56 | 1.03 |
| 12 | Met | CE | 3.02 | 1.14 |
| 12 | Met | CE | 2.27 | 1.11 |
| 17 | Leu | CD2 | 2.75 | 0.99 |
| 17 | Leu | CD1 | 2.52 | 0.90 |
| 25 | Leu | CD2 | 3.61 | 0.87 |
| 25 | Leu | CD2 | 3.04 | 1.12 |
| 25 | Leu | CD1 | 2.31 | 1.03 |
| 29 | Val | CG1 | 3.91 | 0.99 |
| 29 | Val | CG1 | 2.92 | 1.40 |
| 29 | Val | CG2 | 2.42 | 1.06 |
| 31 | Ala | CB | 2.64 | 0.91 |
| 31 | Ala | CB | 2.47 | 1.05 |
| 31 | Ala | CB | 2.53 | 0.87 |
| 32 | Ala | CB | 2.63 | 0.90 |
| 40 | Thr | CG2 | 2.99 | 1.28 |
| 42 | Ala | CB | 2.71 | 0.94 |
| 51 | Thr | CG2 | 3.29 | 1.18 |
| 51 | Thr | CG2 | 2.31 | 1.12 |
| 55 | Ile | CD1 | 2.71 | 1.12 |
| 55 | Ile | CD1 | 2.30 | 1.20 |
| 55 | Ile | CG2 | 2.54 | 0.97 |
| 56 | Leu | CD1 | 3.20 | 1.35 |
| 56 | Leu | CD1 | 2.52 | 1.06 |
| 56 | Leu | CD1 | 2.56 | 1.31 |
| 56 | Leu | CD2 | 2.67 | 0.99 |
| 56 | Leu | CD2 | 2.47 | 1.14 |
| 58 | Ile | CD1 | 2.82 | 1.09 |
| 58 | Ile | CD1 | 2.54 | 1.09 |
| 58 | Ile | CG2 | 3.20 | 1.26 |
| 58 | Ile | CG2 | 3.00 | 0.87 |
| 69 | Thr | CG2 | 2.05 | 1.21 |
| 75 | Leu | CD1 | 3.45 | 1.08 |
| 75 | Leu | CD1 | 2.35 | 0.89 |
| 75 | Leu | CD2 | 2.27 | 0.95 |
| 78 | Ile | CG2 | 3.55 | 1.27 |
| 78 | Ile | CG2 | 2.34 | 1.25 |
| 82 | Ala | CB | 3.39 | 0.89 |
| 82 | Ala | CB | 3.12 | 1.04 |
| 83 | Leu | CD2 | 2.34 | 1.36 |
| 84 | Leu | <u>CD2</u> | 3.58 | 1.18 |
| 84 | Leu | <u>CD1</u> | 2.80 | 1.02 |
| 88 | Ile | CD1 | 3.97 | 1.27 |
| 88 | Ile | CD1 | 2.12 | 1.36 |
| 88 | Ile | CG2 | 2.43 | 1.09 |
| 88 | Ile | CG2 | 2.21 | 1.16 |
| 90 | Ala | CB | 4.32 | 1.13 |
| 92 | Val | CG2 | 3.67 | 0.92 |
| 92 | Val | CG2 | 3.34 | 0.81 |
| 92 | Val | CG2 | 3.08 | 1.02 |
| 95 | Ala | CB | 2.63 | 1.02 |
| 95 | Ala | CB | 2.19 | 0.96 |
| 98 | Ile | CD1 | 3.03 | 1.19 |
| 98 | Ile | CD1 | 2.15 | 0.79 |
| 99 | Val | CG2 | 2.93 | 1.13 |
| 99 | Val | CG2 | 2.79 | 1.07 |
| 105 | Met | CE | 3.46 | 1.16 |
| 107 | Ala | CB | 2.74 | 1.32 |
| 110 | Ala | CB | 2.43 | 1.39 |
| 110 | Ala | CB | 2.03 | 0.96 |
| 118 | Thr | CG2 | 2.09 | 1.03 |
| 120 | Val | CG2 | 3.01 | 1.08 |
| 120 | Val | CG2 | 2.09 | 0.76 |
| 120 | Val | CG1 | 2.42 | 1.59 |
| 122 | Ala | CB | 2.64 | 1.17 |
| 122 | Ala | CB | 2.03 | 1.14 |
| 124 | Ile | CG2 | 2.79 | 0.98 |
| 124 | Ile | CG2 | 2.67 | 0.65 |

**Supplementary Table 8.** Hydrogen bond distances for N-H.

| Residue | Name | Atom | Diff. peak $\sigma$ | X-H (Å) |
| --- | --- | --- | --- | --- |
| 4 | Gly | N | 3.10 | 0.94 |
| 6 | Cys | N | 2.34 | 1.24 |
| 7 | Glu | N | 2.72 | 0.87 |
| 14 | Arg | NE1 | 2.47 | 1.19 |
| 18 | Asp | N | 2.90 | 0.91 |
| 21 | Arg | N | 3.12 | 1.26 |
| 22 | Gly | N | 3.94 | 0.80 |
| 23 | Tyr | N | 4.43 | 1.13 |
| 25 | Leu | N | 2.62 | 0.93 |
| 26 | Gly | N | 3.30 | 1.00 |
| 37 | Asn | N | 2.66 | 1.09 |
| 38 | Phe | N | 2.98 | 1.18 |
| 39 | Asn | N | 3.20 | 1.14 |
| 45 | Arg | N | 2.35 | 0.75 |
| 49 | Gly | N | 2.70 | 0.98 |
| 55 | Ile | N | 2.70 | 0.88 |
| 56 | Ile | N | 2.72 | 1.21 |
| 58 | Ile | N | 3.27 | 1.08 |
| 60 | Ser | N | 3.37 | 0.91 |
| 61 | Arg | N | 4.43 | 0.94 |
| 62 | Trp | N | 2.96 | 0.81 |
| 62 | Trp | NE1 | 2.35 | 1.11 |
| 71 | Gly | N | 2.21 | 1.12 |
| 72 | Ser | N | 2.27 | 1.07 |
| 74 | Asn | N | 2.06 | 0.73 |
| 76 | Cys | N | 2.76 | 1.19 |
| 78 | Ile | N | 2.58 | 0.85 |
| 80 | Cys | N | 2.11 | 0.98 |
| 86 | Ser | N | 2.12 | 1.12 |
| 88 | Ile | N | 2.09 | 0.91 |
| 89 | Thr | N | 4.00 | 1.14 |
| 90 | Ala | N | 3.27 | 1.13 |
| 91 | Ser | N | 2.14 | 0.99 |
| 104 | Gly | N | 2.87 | 1.24 |
| 105 | Met | N | 2.64 | 1.25 |
| 107 | Ala | N | 3.15 | 0.71 |
| 108 | Trp | NE1 | 2.37 | 1.14 |
| 109 | Val | N | 3.68 | 0.81 |
| 111 | Trp | N | 3.22 | 1.21 |
| 118 | Thr | N | 4.54 | 1.02 |
| 119 | Asp | N | 2.50 | 1.26 |
| 120 | Val | N | 3.86 | 1.16 |
| 121 | Asn | N | 2.50 | 0.88 |
| 124 | Ile | N | 3.53 | 0.84 |

**Supplementary Table 9.** Hydrogen bond distances for N-H_2_.

| Residue | Name | Atom | Diff. peak $\sigma$ | X-H (Å) |
| --- | --- | --- | --- | --- |
| 5 | Arg | NH1 | 2.43 | 1.14 |
| 14 | Arg | NH2 | 2.96 | 0.76 |
| 14 | Arg | NH1 | 2.93 | 1.26 |
| 19 | Asn | ND2 | 2.63 | 1.31 |
| 19 | Asn | ND2 | 2.37 | 1.14 |
| 27 | Asn | ND2 | 3.07 | 0.87 |
| 27 | Asn | ND2 | 2.00 | 1.13 |
| 39 | Asn | ND2 | 2.18 | 1.17 |
| 57 | Gln | NE2 | 2.80 | 1.00 |
| 59 | Asn | ND2 | 2.21 | 0.75 |
| 62 | Arg | NH1 | 2.42 | 1.10 |
| 74 | Asn | ND2 | 3.40 | 1.50 |
| 114 | Arg | NH1 | 2.07 | 0.92 |

**Supplementary Table 10.** Hydrogen bond distances for N-H_3_.

| Residue | Name | Atom | Diff. peak $\sigma$ | X-H (Å) |
| --- | --- | --- | --- | --- |
| 1 | Lys | NZ | 3.32 | 1.24 |
| 33 | Lys | NZ | 2.70 | 1.13 |
| 97 | Lys | NZ | 2.96 | 1.02 |

## References

Clabbers, M. T. B., Fisher, S. Z., Coinçon, M., Zou, X. & Xu, H. (2020). Commun. Biol. 3, 417.

Clabbers, M. T. B., Gruene, T., van Genderen, E. & Abrahams, J. P. (2019). Acta Crystallogr. Sect. Found. Adv. 75, 82–93.

Clabbers, M. T. B., Holmes, S., Muusse, T. W., Vajjhala, P. R., Thygesen, S. J., Malde, A. K., Hunter, D. J. B., Croll, T. I., Flueckiger, L., Nanson, J. D., Rahaman, Md. H., Aquila, A., Hunter, M. S., Liang, M., Yoon, C. H., Zhao, J., Zatsepin, N. A., Abbey, B., Sierecki, E., Gambin, Y., Stacey, K. J., Darmanin, C., Kobe, B., Xu, H. & Ve, T. (2021). Nat. Commun. 12, 2578.

Cowley, J. M. (1995). Diffraction physics. Amsterdam: North-Holland.

Cowtan, K. (2006). Acta Crystallogr. D Biol. Crystallogr. 62, 1002–1011.

Dorset, D. L. (1995). Structural electron crystallography. New York: Plenum Press.

Emsley, P., Lohkamp, B., Scott, W. G. & Cowtan, K. (2010). Acta Crystallogr. D Biol. Crystallogr. 66, 486–501.

Eriksson, U. K., Fischer, G., Friemann, R., Enkavi, G., Tajkhorshid, E. & Neutze, R. (2013). Science.

Evans, P. R. & Murshudov, G. N. (2013). Acta Crystallogr. D Biol. Crystallogr. 69, 1204–1214.

Foadi, J., Woolfson, M. M., Dodson, E. J., Wilson, K. S., Jia-xing, Y. & Chao-de, Z. (2000). Acta Crystallogr. D Biol. Crystallogr. 56, 1137–1147.

Fujiwara, K. (1959). J. Phys. Soc. Jpn. 14, 1513–1524.

Gonen, T., Cheng, Y., Sliz, P., Hiroaki, Y., Fujiyoshi, Y., Harrison, S. C. & Walz, T. (2005). Nature. 438, 633–638.

Gruene, T., Hahn, H. W., Luebben, A. V., Meilleur, F. & Sheldrick, G. M. (2014). J. Appl. Crystallogr. 47, 462–466.

Gruene, T., Wennmacher, J. T. C., Zaubitzer, C., Holstein, J. J., Heidler, J., Fecteau-Lefebvre, A., De Carlo, S., Müller, E., Goldie, K. N., Regeni, I., Li, T., Santiso-Quinones, G., Steinfeld, G., Handschin, S., van Genderen, E., van Bokhoven, J. A., Clever, G. H. & Pantelic, R. (2018). Angew. Chem. Int. Ed. 57, 16313–16317.

Guo, H., Franken, E., Deng, Y., Benlekbir, S., Singla Lezcano, G., Janssen, B., Yu, L., Ripstein, Z. A., Tan, Y. Z. & Rubinstein, J. L. (2020). IUCrJ. 7, 860–869.

Hattne, J., Shi, D., Glynn, C., Zee, C.-T., Gallagher-Jones, M., Martynowycz, M. W., Rodriguez, J. A. & Gonen, T. (2018). Structure. 26, 759–766.e4.

Henderson, R. (1995). Q. Rev. Biophys. 28, 171–193.

Howard, E. I., Sanishvili, R., Cachau, R. E., Mitschler, A., Chevrier, B., Barth, P., Lamour, V., Van Zandt, M., Sibley, E., Bon, C., Moras, D., Schneider, T. R., Joachimiak, A. & Podjarny, A. (2004). Proteins Struct. Funct. Bioinforma. 55, 792–804.

Jones, C. G., Martynowycz, M. W., Hattne, J., Fulton, T. J., Stoltz, B. M., Rodriguez, J. A., Nelson, H. M. & Gonen, T. (2018). ACS Cent. Sci. 4, 1587–1592.

Kabsch, W. (2010). Acta Crystallogr. D Biol. Crystallogr. 66, 125–132.

Kimura, Y., Vassylyev, D. G., Miyazawa, A., Kidera, A., Matsushima, M., Mitsuoka, K., Murata, K., Hirai, T. & Fujiyoshi, Y. (1997). Nature. 389, 206–211.

Latychevskaia, T. & Abrahams, J. P. (2019). Acta Crystallogr. Sect. B Struct. Sci. Cryst. Eng. Mater. 75, 523–531.

Leapman, R. D. & Sun, S. (1995). Ultramicroscopy. 59, 71–79.

Liu, S. & Gonen, T. (2018). Commun. Biol. 1, 38.

Maki-Yonekura, S., Kawakami, K., Hamaguchi, T., Takaba, K. & Yonekura, K. (2021). Hydrogen properties and charges in a sub-1.2 Å resolution cryo-EM structure revealed by a cold field emission beam Biophysics.

Martynowycz, M. W., Clabbers, M. T. B., Hattne, J. & Gonen, T. (2021). Ab initio phasing macromolecular structures using electron-counted MicroED data Biochemistry.

Martynowycz, M. W., Clabbers, M. T. B., Unge, J., Hattne, J. & Gonen, T. (2021). Proc. Natl. Acad. Sci. 118, e2108884118.

Martynowycz, M. W., Khan, F., Hattne, J., Abramson, J. & Gonen, T. (2020). Proc. Natl. Acad. Sci. 117, 32380–32385.

Martynowycz, M. W., Shiriaeva, A., Ge, X., Hattne, J., Nannenga, B. L., Cherezov, V. & Gonen, T. (2021). Proc. Natl. Acad. Sci. 118, e2106041118.

McCoy, A. J., Grosse-Kunstleve, R. W., Adams, P. D., Winn, M. D., Storoni, L. C. & Read, R. J. (2007). J. Appl. Crystallogr. 40, 658–674.

Mitsuoka, K., Hirai, T., Murata, K., Miyazawa, A., Kidera, A., Kimura, Y. & Fujiyoshi, Y. (1999). J. Mol. Biol. 286, 861–882.

Murshudov, G. N., Skubák, P., Lebedev, A. A., Pannu, N. S., Steiner, R. A., Nicholls, R. A., Winn, M. D., Long, F. & Vagin, A. A. (2011). Acta Crystallogr. D Biol. Crystallogr. 67, 355–367.

Nakane, T., Kotecha, A., Sente, A., McMullan, G., Masiulis, S., Brown, P. M. G. E., Grigoras, I. T., Malinauskaite, L., Malinauskas, T., Miehling, J., Uchański. T., Yu, L., Karia, D., Pechnikova, E. V., de Jong, E., Keizer, J., Bischoff, M., McCormack, J., Tiemeijer, P., Hardwick, S. W., Chirgadze, D. Y., Murshudov, G., Aricescu, A. R. & Scheres, S. H. W. (2020). Nature. 587, 152–156.

Nannenga, B. L., Shi, D., Hattne, J., Reyes, F. E. & Gonen, T. (2014). ELife. 3, e03600.

Nannenga, B. L., Shi, D., Leslie, A. G. W. & Gonen, T. (2014). Nat. Methods. 11, 927–930.

Ogata, H., Nishikawa, K. & Lubitz, W. (2015). Nature. 520, 571–574.

Palatinus, L., Brázda, P., Boullay, P., Perez, O., Klementová, M., Petit, S., Eigner, V., Zaarour, M. & Mintova, S. (2017). Science. 355, 166–169.

Purdy, M. D., Shi, D., Chrustowicz, J., Hattne, J., Gonen, T. & Yeager, M. (2018). Proc. Natl. Acad. Sci. 115, 13258–13263.

Rodriguez, J. A., Ivanova, M. I., Sawaya, M. R., Cascio, D., Reyes, F. E., Shi, D., Sangwan, S., Guenther, E. L., Johnson, L. M., Zhang, M., Jiang, L., Arbing, M. A., Nannenga, B. L., Hattne, J., Whitelegge, J., Brewster, A. S., Messerschmidt, M., Boutet, S., Sauter, N. K., Gonen, T. & Eisenberg, D. S. (2015). Nature. 525, 486–490.

Sawaya, M. R., Rodriguez, J., Cascio, D., Collazo, M. J., Shi, D., Reyes, F. E., Hattne, J., Gonen, T. & Eisenberg, D. S. (2016). Proc. Natl. Acad. Sci. 113, 11232–11236.

Takaba, K., Maki-Yonekura, S., Inoue, I., Tono, K., Hamaguchi, T., Kawakami, K., Naitow, H., Ishikawa, T., Yabashi, M. & Yonekura, K. (2021). Hydrogen properties in an organic molecule revealed by XFEL and electron crystallography Chemistry.

Walsh, M. A., Schneider, T. R., Sieker, L. C., Dauter, Z., Lamzin, V. S. & Wilson, K. S. (1998). Acta Crystallogr. D Biol. Crystallogr. 54, 522–546.

Wang, J., Dauter, M., Alkire, R., Joachimiak, A. & Dauter, Z. (2007). Acta Crystallogr. D Biol. Crystallogr. 63, 1254–1268.

Williams, C. J., Headd, J. J., Moriarty, N. W., Prisant, M. G., Videau, L. L., Deis, L. N., Verma, V., Keedy, D. A., Hintze, B. J., Chen, V. B., Jain, S., Lewis, S. M., Arendall, W. B., Snoeyink, J., Adams, P. D., Lovell, S. C., Richardson, J. S. & Richardson, D. C. (2018). Protein Sci. 27, 293–315.

Winn, M. D., Ballard, C. C., Cowtan, K. D., Dodson, E. J., Emsley, P., Evans, P. R., Keegan, R. M., Krissinel, E. B., Leslie, A. G. W., McCoy, A., McNicholas, S. J., Murshudov, G. N., Pannu, N. S., Potterton, E. A., Powell, H. R., Read, R. J., Vagin, A. & Wilson, K. S. (2011). Acta Crystallogr. D Biol. Crystallogr. 67, 235–242.

Xu, H., Lebrette, H., Clabbers, M. T. B., Zhao, J., Griese, J. J., Zou, X. & Högbom, M. (2019). Sci. Adv. 5, eaax4621.

Yamashita, K., Palmer, C. M., Burnley, T. & Murshudov, G. N. (2021). Acta Crystallogr. Sect. Struct. Biol. 77, 1282–1291.

Yonekura, K., Ishikawa, T. & Maki-Yonekura, S. (2019). J. Struct. Biol. 206, 243–253.

Yonekura, K., Kato, K., Ogasawara, M., Tomita, M. & Toyoshima, C. (2015). Proc. Natl. Acad. Sci. 112, 3368–3373.

Yonekura, K., Matsuoka, R., Yamashita, Y., Yamane, T., Ikeguchi, M., Kidera, A. & Maki-Yonekura, S. (2018). IUCrJ. 5, 348–353.

